# *Pseudomonas* mRNA 2.0: Boosting Gene Expression Through Enhanced mRNA Stability and Translational Efficiency

**DOI:** 10.1101/787127

**Authors:** Dário Neves, Stefan Vos, Lars M. Blank, Birgitta E. Ebert

## Abstract

High gene expression of enzymes partaking in recombinant production pathways is a desirable trait among cell factories belonging to all different kingdoms of life. High enzyme abundance is generally aimed for by utilizing strong promoters, which ramp up gene transcription and mRNA levels. Increased protein abundance can alternatively be achieved by optimizing the expression on the post-transcriptional level. Here, we evaluated protein synthesis with a previously proposed optimized gene expression architecture, in which mRNA stability and translation initiation are modulated by genetic parts such as self-cleaving ribozymes and a bicistronic design, which have initially been described to support the standardization of gene expression. The optimized gene expression architecture was tested in *Pseudomonas taiwanensis* VLB120, a promising, novel microbial cell factory. The expression cassette was employed on a plasmid basis and after single genomic integration. We used three constitutive and two inducible promoters to drive the expression of two fluorescent reporter proteins and a short acetoin biosynthesis pathway. The performance was confronted with that of a traditional expression cassette harboring the same promoter and gene of interest but lacking the genetic parts for increased expression efficiency. The optimized expression cassette granted higher protein abundance independently of the expression basis or promoter used proving its value for applications requiring high protein abundance.

## 1 Introduction

Cell factories have become an established role player in the sustainable production of chemicals and biological products proven with hundreds of billions of USD/year value on global markets (Davy et al., 2017). A commonality in the development of such cell factories is the continuous pursuit of increased productivities through directed or selection-based genetic engineering methods. With both approaches, increasing activity of the partaking pathways commonly leads to the desired rise in productivity. High enzyme activity can be achieved by optimization of transcription, translation, post-translational modifications, and the process conditions (Liu et al., 2013). A common strategy is to employ strong promoters to overexpress product biosynthesis genes. Highly-active promoters achieve increased protein production rates by increasing the respective mRNA levels in the cell. However, previous studies have shown that high, recombinant gene expression leads to metabolic burden and consequently to growth impairment (Borkowski et al., 2016; Carneiro et al., 2013). Such hindrances are related to the drainage of biosynthetic precursors, such as nucleotides, or seizing of the cellular transcriptional machinery.

In recent years, Synthetic Biology parts emerged that support high enzyme activities without the need for strong gene expression, thereby contributing to diminishing competition and depletion of the cellular mRNA pool and lightening the metabolic burden. Two translation-focused approaches can be distinguished that target to optimize translation rather than transcription. To this end, the first approach attempts to stabilize mRNA, whereas the second seeks to increase translational efficiency. Increasing mRNA stability is possible by placing stabilizing sequences in the 5’ untranslated region (UTR) that avoid endoribonuclease attacks through their secondary structures as in *Escherichia coli* (Carrier and Keasling, 1997; Viegas et al., 2018). The implementation of ribozymes upstream of the ribosome binding site (RBS) allows the insulation of the desired expression cassette from the genetic context (Lou et al., 2012). Besides the intrinsic cleaving activity, the ribozymes developed by Lou et al. contained a 23 nucleotide hairpin downstream of the catalytic core, which additionally adds an mRNA stabilizing trait to this genetic part (Clifton et al., 2018). One approach within the second category, which focuses on translational efficiency, allows increased expression levels by facilitating the access of the ribosome to the RBS. In the traditional operon architecture (Figure 1 A), it is possible that gene of interest (GOI)-dependent secondary structures arise. This folding of the mRNA can block the access of ribosomes to the RBS thereby compromising the desired gene expression (Salis et al., 2009). Mutalik et al. developed a ‘bicistronic design’ which takes advantage of the intrinsic helicase activity of ribosomes to unveil any RBS-GOI dependent secondary structure and consequently achieve GOI-independent expression (Mutalik et al., 2013). In the bicistronic design, a short leading peptide cistron is allocated upstream of the GOI. The RBS of the GOI is enclosed within the coding sequence of the leading cistron, whereas the start codon of the GOI is fused to the stop codon of the leading cistron. The leading RBS-small peptide combination is known to not create any secondary structures, which assures the binding of a ribosome. Once the ribosome binds to the first RBS and starts to translate the leading peptide, any possible downstream RBS-GOI dependent secondary structures are unveiled by its intrinsic helicase activity exposing the RBS of the GOI. Otto et al. recently combined the bicistronic design with an upstream ribozyme to increase the translational efficiency of heterologous genes integrated in rRNA operons. Here the intention of the ribozyme integration was not to stabilize the mRNA but to increase translation efficiency by reducing a potential steric hindrance by the bulky 5’ 16S mRNA flank and thereby facilitating ribosome docking. To this end, the ribozyme was placed upstream of the RBS, and this integration indeed resulted in a substantial increase in protein production (Otto et al., 2019).

**Figure 1.**
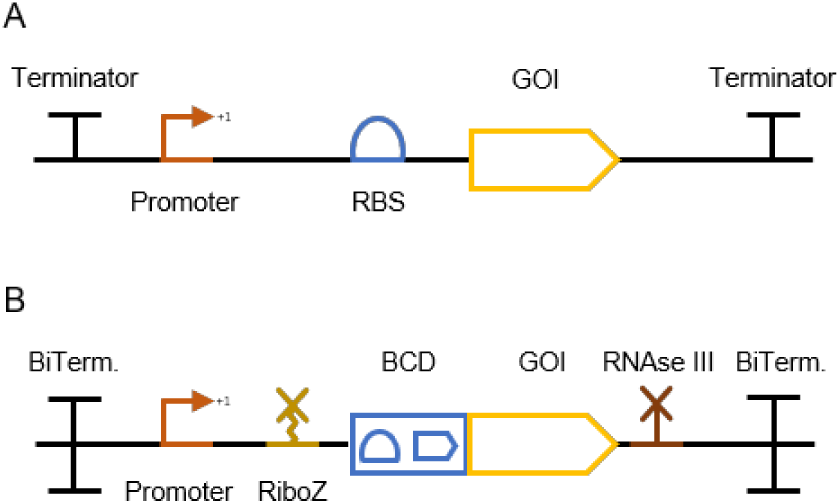
Gene expression cassette architectures represented with glyphs compliant with Synthetic Biology Open Language Visual (SBOLv). (A) A traditional gene expression cassette comprising a promoter, an RBS, and a gene of interest (GOI), (B) an optimized gene expression cassette as proposed by Nielsen et al., framed between two bidirectional terminators (Bi. term.) and encompassing a promoter, a ribozyme (RiboZ), the bicistronic design (BCD) developed by Mutalik et al.(Mutalik et al., 2013), a GOI and an RNAse III site.

Nielsen et al. proposed the compilation of these and further genetic parts into an overall standardized gene expression cassette for the assembly of genetic circuits (Figure 1 B) (Nielsen et al., 2013). Besides the incorporation of the gene expression parts described above, Nielsen et al. proposed the isolation of the expression cassette with bidirectional terminators on both ends and the integration of an RNAse III site downstream of the GOI to further reduce context-specific effects. The impact of an RNAse III site downstream of the GOI was evaluated by Cambray et al. within the scope of reliable terminator characterization (Cambray et al., 2013). Their extensive work showed that despite an overall decrease in expression level upon RNA III site incorporation, possibly due to a shorter mRNA half-life, a much narrower and precise expression level can be achieved. These promoter-independent gene expression tools have been individually characterized but, to our knowledge, a possible synergistic and expression enhancing effect of their combination is yet to be explored.

In this work we constructed optimized gene expression cassettes based on the architecture proposed by Nielsen et al. and evaluated the performance against traditional configurations using two fluorescence proteins (msfGFP and mCherry) and recombinant acetoin production as readout (Landgraf, 2012). The constructed, optimized gene expression cassettes were evaluated on a plasmid basis and after single-copy genomic integration (Wierckx et al., 2005). Overall, the traditional and optimized gene expression cassette variants were characterized with three constitutive and two inducible promoters. Reducing the overall size of the optimized gene expression cassette while maintaining its performance was also targeted in this work. Besides the characterization of several constructs, qPCR analysis was performed to elucidate the role of mRNA stability in altered protein expression. We chose *Pseudomonas taiwanensis* VLB120 as expression host as this Gram-negative bacterium exhibits industrial relevant, metabolic capabilities such as broad carbon source utilization, the ability to proliferate in the presence of organic solvents, and an almost byproduct free metabolism (Köhler et al., 2013; Park et al., 2007). *P. taiwanensis* VLB120 has been proven a suitable biocatalyst for the production of (*S*)-styrene oxide, phenol, isobutyric acid, and 4-hydroxybenzoic acid (Lang et al., 2014; Lenzen et al., 2019; Panke et al., 1998; Wynands et al., 2018). The novel expression device developed in this study contributes to more effective engineering of this emergent and promising biocatalysts and other prokaryotic cell factories

## 2 Materials and Methods

### 2.1 Media and Growth Conditions

Liquid cultures were grown in a horizontal rotary shaker with a shaking frequency of 200 rpm and a throw of 50 mm in LB medium or LB medium supplemented with 5 g/L glucose and buffered with 11.64 g/L K_2_HPO_4_ and 4.89 g/L NaH_2_PO_4_ (LB_mod_). *Pseudomonas* strains were grown at 30°C, whereas *E. coli* was grown at 37°C. Solid LB was prepared by adding 1.5 % (w/v) agar to the medium. Antibiotics were supplemented to the medium for plasmid maintenance and selection purposes. Kanamycin sulfate was added at a concentration of 50 mg/L for *Pseudomonas* and *E. coli*. Gentamycin was used at a concentration of 25 mg/L for both species. Tetracycline was added only to solid media at a concentration of 30 mg/L for *Pseudomonas* and 10 mg/L for *E. coli*. To induce the pTN1 derived plasmids harboring the *nag*R*/*P*nagAa* promoter system controlling the expression of the fluorescent proteins or the acetoin pathway, 0.01 mM or 1 mM of sodium salicylate was added, respectively. The inducer isopropyl-β-D-1 thiogalactopyranoside (IPTG) was used at a concentration of 1 mM to induce the P_trc_ controlled constructs integrated into the *att*Tn7 site of *Pseudomonas* strains.

The acetoin producing strains were cultivated in airtight 500 mL serum flasks containing 50 mL of LB_mod_ supplemented with gentamycin (see above). The main cultures were inoculated from an overnight pre-culture to an OD_600_ of 0.1. The plasmid-based acetoin pathway genes were induced with sodium salicylate once the cultures reached an OD_600_ of 1. Afterward, samples were collected for HPLC analysis.

The chemicals used were purchased from Merck (Darmstadt, Germany), Sigma-Aldrich (St. Louis, MO, USA) or Carl Roth (Karlsruhe, Germany) unless stated otherwise. Pharmaceutical grade glycerol was kindly provided by Bioeton (Kyritz, Germany).

### 2.2 Plasmid and strain construction

All plasmids were constructed through Gibson assembly (Gibson et al., 2009) using the NEBuilder HiFi DNA Assembly kit (New England Biolabs, Ipswich, MA, USA). Primers used in this study were purchased from Eurofins Genomics (Ebersberg, Germany) as unmodified DNA oligonucleotides. PCR amplification of DNA for cloning purposes was performed using the Q5 High-Fidelity Polymerase (New England Biolabs, Ipswich, MA, USA). All primers and plasmids are listed in the Supplementary Information Table S1. The genes *ilvB* (from *E. coli* K-12 MG1655, Uniprot P08142, with C83S mutation for improved Kcat/Km) and *aldB* (from *B. brevis*, Uniprot P23616) were codon-optimized for *P. taiwanensis* VLB120 using the online tool OPTIMIZER (Puigbo et al., 2007). Settings were as follows: genetic code, eubacterial; method, guided random; undesired restriction sites were manually excluded, and rare codons with less than 6% usage were avoided by manipulating the input codon usage table. Both codon-optimized genes and their corresponding optimized gene expression parts were ordered as synthetic DNA fragments from Thermo Fisher Scientific; the sequences can be found in the Supplementary Information A. The assembled plasmids were transformed into either NEB® 5-alpha chemically competent *E. coli* (New England Biolabs, Ipswich, MA, USA) or One Shot™ PIR2 Chemically Competent *E. coli* (Thermo Fisher Scientific) cells through heat shock according to the supplier’s protocol. Assembled plasmids were transformed into *P. taiwanensis* VLB120 by electroporation using a GenePulser Xcell (BioRad, Hercules, CA, USA) (settings: 2 mm cuvette gap, 2.5 kV, 200 Ω,25 μF). For DNA integration into the *att*Tn7 locus, the mini-Tn7 delivery vector backbone developed by Zobel et al. (Zobel et al., 2015). was used and deployed through mating procedures. For mating events, the *E. coli* donor harboring the mini-Tn7 vector with the constructs to be integrated, the helper strain *E. coli* HB101 pRK2013, the *E. coli* DH5 λpir expressing the transpose operon *tnsABCD* and the recipient strain were streaked on top of each other on a LB agar plate and incubated at 30°C for 12-24 h. Further on, cell material was taken from the bacterial lawn, resuspended in 0.9 % (w/v) sodium chloride solution and plated on selective cetrimide agar plates. *E. coli* and *Pseudomonas* transformants were screened through colony PCR using the OneTaq 2×Master Mix with standard buffer after lysing colony cell material in alkaline polyethylene glycol, as described by Chomczynski and Rymaszewski.(Chomczynski and Rymaszewski, 2006) Successful plasmid constructions and genome integrations were confirmed by Sanger sequencing performed by Eurofins Genomics. All strains and primers used in this work are shown in Table 1.

**Table 1.**
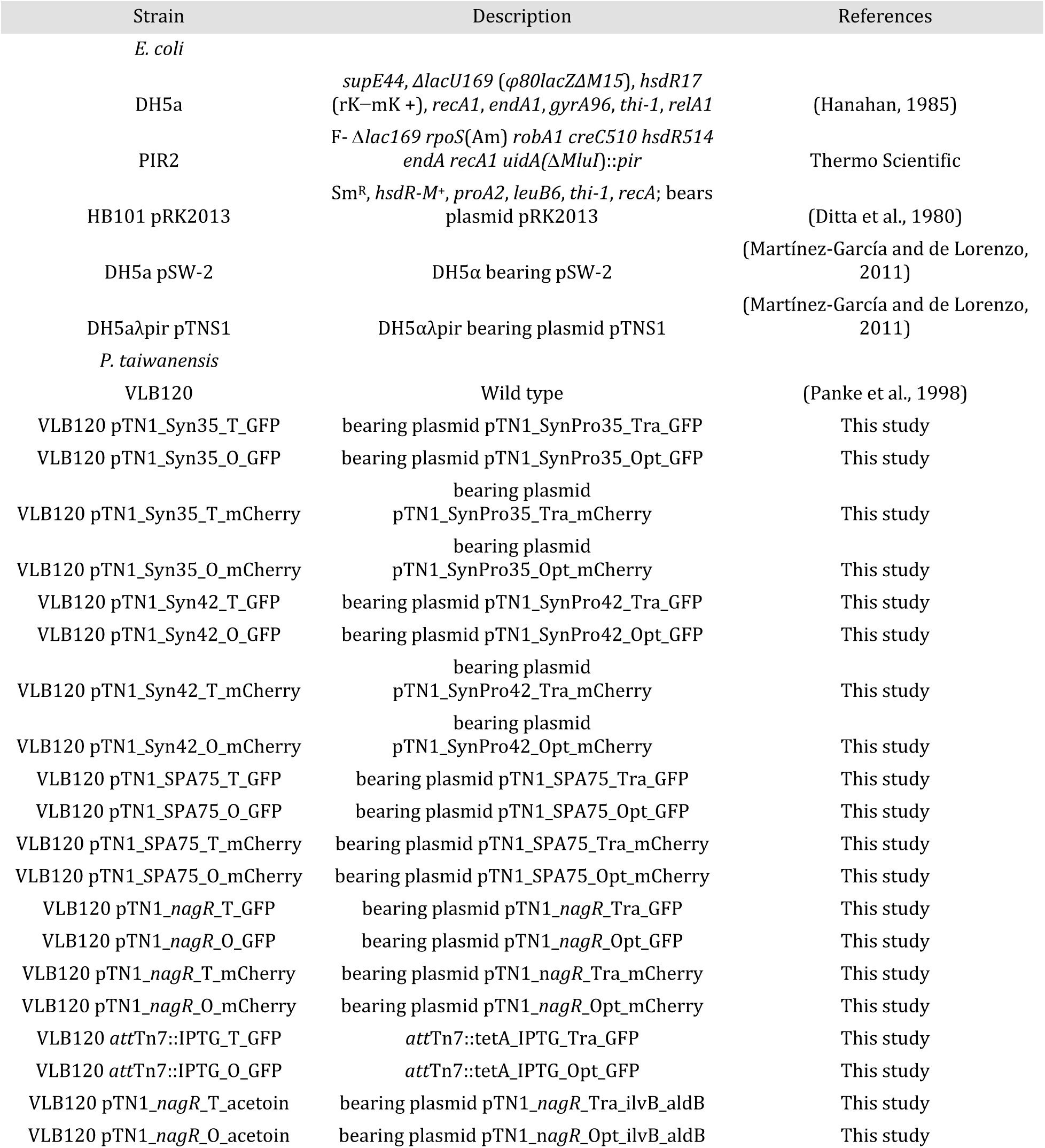
Strains used in this study.

### 2.3 Gravimetric cell dry weight determination

Overnight cultures of *P. taiwanensis* VLB120 were diluted to pre-established optical densities and filtered through membrane filters with a pore size of 0.2 µM which were previously weighted (w_0_) after being dried in a microwave for 3 min at 350 W. The filter cake was washed three times with distilled water, dried in the microwave for 8 min at 350 W and cooled down in a desiccator before weighing (w_1_). The cell dry weight (CDW) was calculated by subtracting w_0_ from w_1_ and divided by the volume of the filtered suspensions. 190 µL of the same cell suspensions were transferred to a 96-well microtiter plate (Greiner Bio One), and scattered light signals were recorded at 620 nm with a gain of 30. Linear regression between the CDW and scattered light signals of the pre-established cell suspensions retrieved the conversion factor between these two units.

### 2.4 Fluorescent measurements

All *Pseudomonas* strains expressing fluorescent reporter proteins were characterized in the microbioreactor system BioLector (m2p-labs, Baesweiler, Germany). Cultivations were performed at 30°C with a shaking frequency of 900 rpm and 85 % humidity in a 96-well microtiter plate (Greiner Bio One) containing 190 µL of LB media supplemented with required antibiotics for plasmid maintenance and sealed with evaporation reducing foil (Greiner Bio One). The main cultures were inoculated from an overnight pre-culture to an OD_600_ of 0.1. Growth was measured through scattered light signal at 620 nm with a gain of 30, msfGFP fluorescence was excited at 485 nm and emission was measured at 520 nm. The measurements were performed with gains of 50 and 70 due to signal overflow of the stronger constructs. mCherry fluorescence was excited at 580 nm, and emission was measured at 610 nm with a gain of 100. Scattered light values were converted into cell dry weight concentrations with a predetermined calibration curve. Inducer was added to the cultures during the early exponential growth phase. To allow comparison of fluorescence values recorded with different gains, the msfGFP signals were converted into µM units of fluorescein, which was dissolved in 100 mM boric acid (See Supplementary information B for the calibration curves). Biological triplicates were performed, and errors presented as the standard deviation of the mean.

### 2.5 qPCR assays for mRNA stability assessment

Biological triplicates from *P. taiwanensis* VLB120 harboring either the plasmid pTN1_SPA75_Tra_GFP or pTN1_SPA75_Opt_GFP were grown overnight in LB_mod_ supplemented with 25 mg/mL gentamycin from a glycerol stock. On the following day, the main cultures with the same medium were inoculated to an initial OD_600_ of 0.1. Once the cultures reached an OD of 1, 1 mg/mL rifampicin and 40 µg/mL nalidixic acid were added simultaneously to cease DNA replication and transcription, respectively. 2 mL samples were retrieved, centrifuged, and cell pellets flash-frozen in liquid nitrogen and stored at -80°C until further sample treatment. Cell pellets were suspended in 800 µL of DNA/RNA protecting buffer from the Monarch Total RNA MiniPrep kit (New England Biolabs, Ipswich, MA, USA), transferred into the ZR S6012-50 Bashing beads lysis tubes (Zymo Research, Irvine, CA, USA), and mechanically disrupted for 1 min. The lysate was transferred into a fresh reaction tube and centrifuged for 2 min at 13,000 rpm. The supernatant was transferred into a fresh tube, and the protocol proceeded as described in the Monarch Total RNA MiniPrep kit. qPCR experiments with *msfGFP* and *rpoB* primers pairs were performed with 1 µL of each RNA sample to confirm that the samples were not contaminated with either genomic or plasmid DNA. 80 ng of RNA of each sample was converted into cDNA using the LunaScript RT SuperMix kit (New England Biolabs, Ipswich, MA, USA). qPCR experiments were performed with the Luna Universal qPCR Master Mix (New England Biolabs, Ipswich, MA, USA). Primer pairs efficiencies for the housekeeping gene *rpoB* and target gene *msfGFP* can be seen in Supplementary Information C. qPCR of samples was performed with 1 µL of the reverse transcription reaction mixtures. The qPCR was performed with the CFX96 Real-Time PCR Detection System (Biorad, Hercules, CA, USA). qPCR reactions were performed in technical triplicates. Absolute amounts of mRNA transcripts of *msfGFP* and *rpoB* were quantified using the linear calibration curves used for primer pairs efficiencies, which were constructed with either the plasmid harboring the traditional *msfGFP* expression cassette under the control of the SPA75 or genomic DNA, respectively. The data was analyzed with the Bio-Rad CFX Manager and Microsoft Excel software. As we normalized the *msfGFP* mRNA abundance data with the transcript abundance of the housekeeping gene *rpoB*, the determination of an absolute decay rate for the *msfGFP* mRNA was not possible. We calculated the delta between the decay rates of the *msfGFP* and *rpoB* mRNA instead. Assuming a first-order degradation kinetic for the mRNA of both genes, the time profile of the normalized mRNA data can be described with Equation 1.

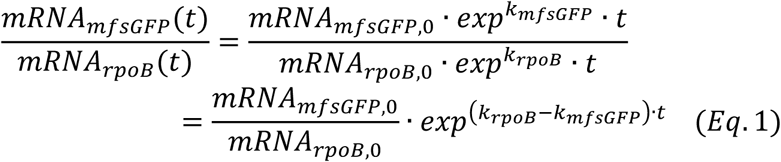

A nonlinear least square algorithm was used in Matlab (The MathWorks Inc., Natick, MA, USA) to fit the experimental data to Equation 1 and to determine the difference in the decay rates of the *rpoB* and the *mfsGFP* gene.

### 2.6 Analytical methods

The samples taken during the acetoin producing cultivations were centrifuged at 13,000 rpm for 1 minute, and the supernatant was stored at -20°C until further analysis. To follow the consumption of glucose and production of acetoin, a Beckman System Gold 126 Solvent Module with an organic acid resin column (Polystyrene divinylbenzene copolymer (PS DVB), 300 × 8.0 mm, CS-Chromatographie) was used with 5 mM H_2_SO_4_ as eluent at a flow of 0.6 mL h^-1^ for 30 minutes at 30°C. Detection was realized with a System Gold 166 UV detector (Beckman Coulter) and a Smartline RI Detector 2300 (KNAUER Wissenschaftliche Geräte, Berlin, Germany).

## 3 Results and Discussion

### 3.1 Characterization of plasmid-based, constitutive fluorescent protein expression

The optimal gene expression profile depends on the specific application. Generally, the use of robust, constitutive promoters is prioritized over inducible promoters for large-scale production as they render the addition of inducers unnecessary and therefore contribute the cost efficiency of microbial fermentations. To evaluate the impact and applicability of the consolidated, optimized expression architecture on this type of promoters, we selected two synthetic promoters, Syn42 and Syn35, created by Zobel et al. (originally referred to as BG42 and BG35, respectively), whereas the third one, SPA75, was obtained from a synthetic promoter library created by Neves and Liebal et al. (manuscript in preparation) (Zobel et al., 2015). The promoters Syn42 and SPA75 possess a rather similar and high expression strength in *P. taiwanensis* VLB120, while the promoter Syn35 exhibits around 25% of the expression strength of Syn42. These promoters were encompassed within the optimized and traditional gene expression cassette in the pTN1 plasmid backbone, a vector used by the *Pseudomonas* scientific community (Figure 2 A) (Schmitz et al., 2015; Verhoef et al., 2009; Wierckx et al., 2005). The optimized gene expression cassette was framed between two bidirectional terminators to uncouple transcription from its genetic context. For this purpose, two bidirectional terminators characterized by Chen et al., ECK120026481 and ECK120011170, were selected to insulate, respectively, the 5’ and 3’ end of the optimized gene expression cassette (Chen et al., 2013). The ribozyme RiboJ, characterized by Lou et al., and the bicistronic design BCD2, developed by Mutalik et al., were placed downstream of the selected promoters (Lou et al., 2012; Mutalik et al., 2013). The last genetic part included was the RNAse III site R1.1, characterized by Cambray et al., and placed downstream of the GOI (Cambray et al., 2013). The traditional versions of the gene expression cassettes were obtained by omitting the enhancing genetic parts and including the 2^nd^ RBS of the BCD2 to maintain the ribosome affinity towards the mRNA between the two expression systems (Figure 2 A).

**Figure 2.**
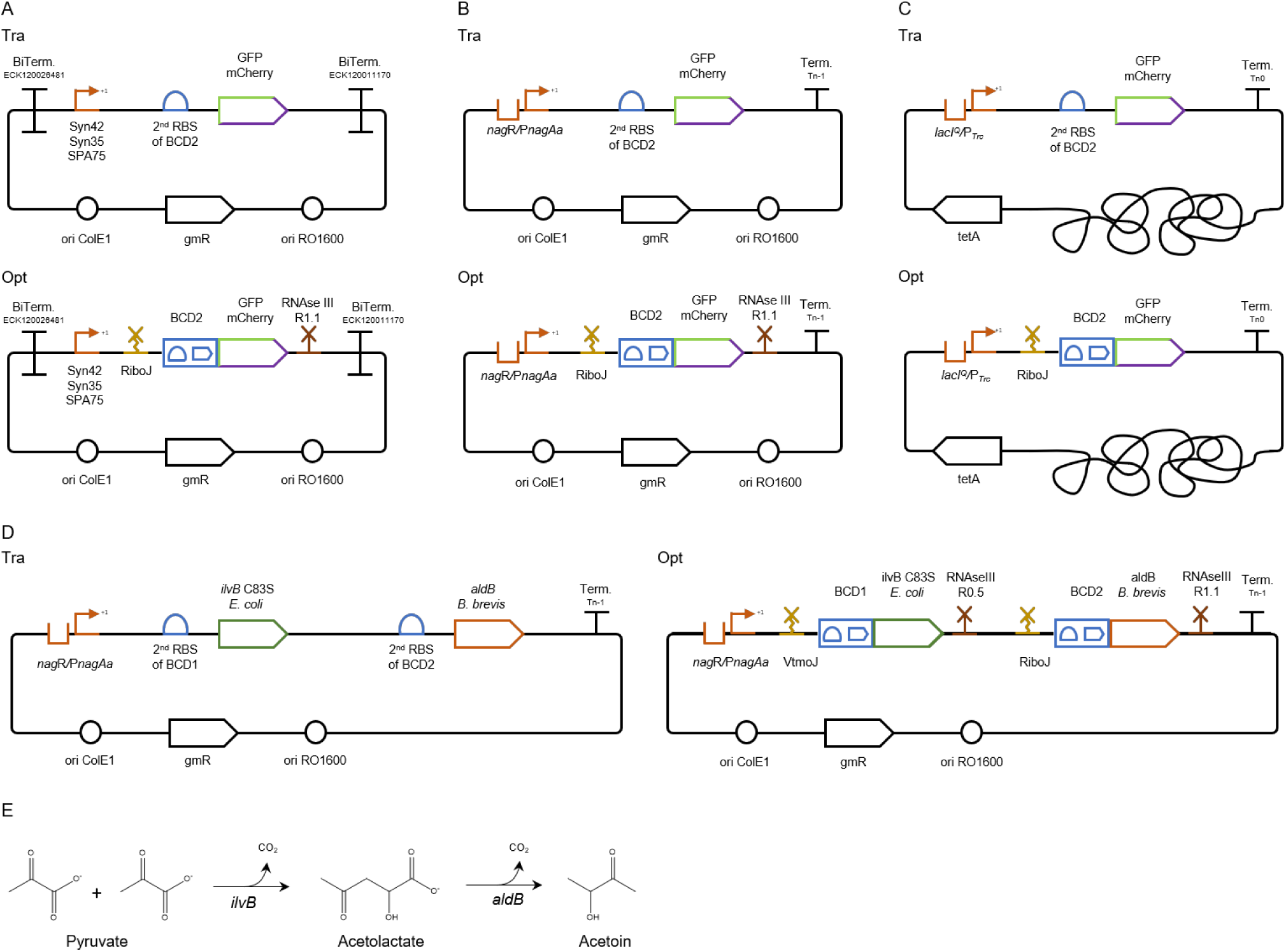
Gene expression constructs evaluated within this work. Traditional and optimized gene expression cassettes for (A) the plasmid-based evaluation of the expression of two fluorescent reporters (msfGFP and mCherry) under the control of three constitutive promoters (Syn42, Syn35, and SPA75) and (B) the salicylate inducible PnagAa promoter; (C) Traditional and optimized gene expression cassettes for single genomic integration into the attTn7 site. The expression with these constructs was evaluated using the fluorescent reporter msfGFP under the control of the IPTG inducible PTrc promoter; (D) traditional and optimized gene expression cassettes for the plasmid-based evaluation of an acetoin pathway under the control of the salicylate inducible PnagAa promoter, (E) acetoin pathway comprising the C83S ilvB mutant from E. coli and aldB from Brevibacillus brevis; Tra, traditional gene expression cassette; Opt, optimized gene expression cassette; RBS, ribosomal binding site; Bi. term., bidirectional terminator; BCD, bicistronic design; VtmoJ, RiboJ, synthetic ribozyme; RNAseIII R1.1 and R0.5, RNase III restriction sites; *gmR*, gentamycin resistance gene; *tetA*, tetracycline resistance gene; ori ColE1 and ori RO1600, origins of replication.

The twelve constructs were evaluated in microtiter plate cultivations with online measurements of fluorescence and scattered light. The scattered light values were converted into cell dry weight units, whereas the arbitrary *msfGFP* fluorescence units were transformed into equivalents of fluorescein (µM) to allow a direct comparison between experiments ran with different measurement settings. The *mCherry* fluorescence values were not converted since all experiments were performed with the same settings. However, we propose the broad implementation of such standardized fluorescence units to facilitate results comparison within the scientific community. Gene expression with the different constructs was characterized by the slope of the linear regression between measured fluorescence and cell dry weight, which indicates a specific expression strength.

All optimized gene expression constructs with the three tested constitutive promoters resulted in a substantial increase in fluorescence compared with their traditional counterparts (Figure 3 A). While the ranking of the promoter strength was maintained with the optimized gene expression cassettes, absolute differences in the level of expression were reduced as a much higher increase was observed for the weaker Syn35 promoter expressing mCherry (Table 2). The lower fold increase in expression strength for the strong promoters suggests that the full potential of the optimized gene expression cassette is not reached here because of other cellular limitations, such as RNA polymerase availability.

**Table 2.** Pairwise fold-changes of specific fluorescence (fluorescence per g cell dry weight) between the optimized and traditional gene expression cassettes calculated as described by Clifton et al (Clifton et al., 2018).

| Expression System |  | pTN1 plasmid |  |  |  | -attTn7:: |
| --- | --- | --- | --- | --- | --- | --- |
| Promoter | Syn35 | Syn42 | SPA75 | *nagR/PnagAa* |  | *lacI<sup>Q</sup>/P<sub>Trc</sub>* |
| msfGFP | 8.7±1.6 | 5.0±0.9 | 3.9±0.3 | 4.1±0.7 |  | 14.7±1.6 |
| mCherry | 11.3±3.8 | 3.8±0.2 | 2.9±0.4 | 3.9±0.7 |  | - |

**Figure 3.**
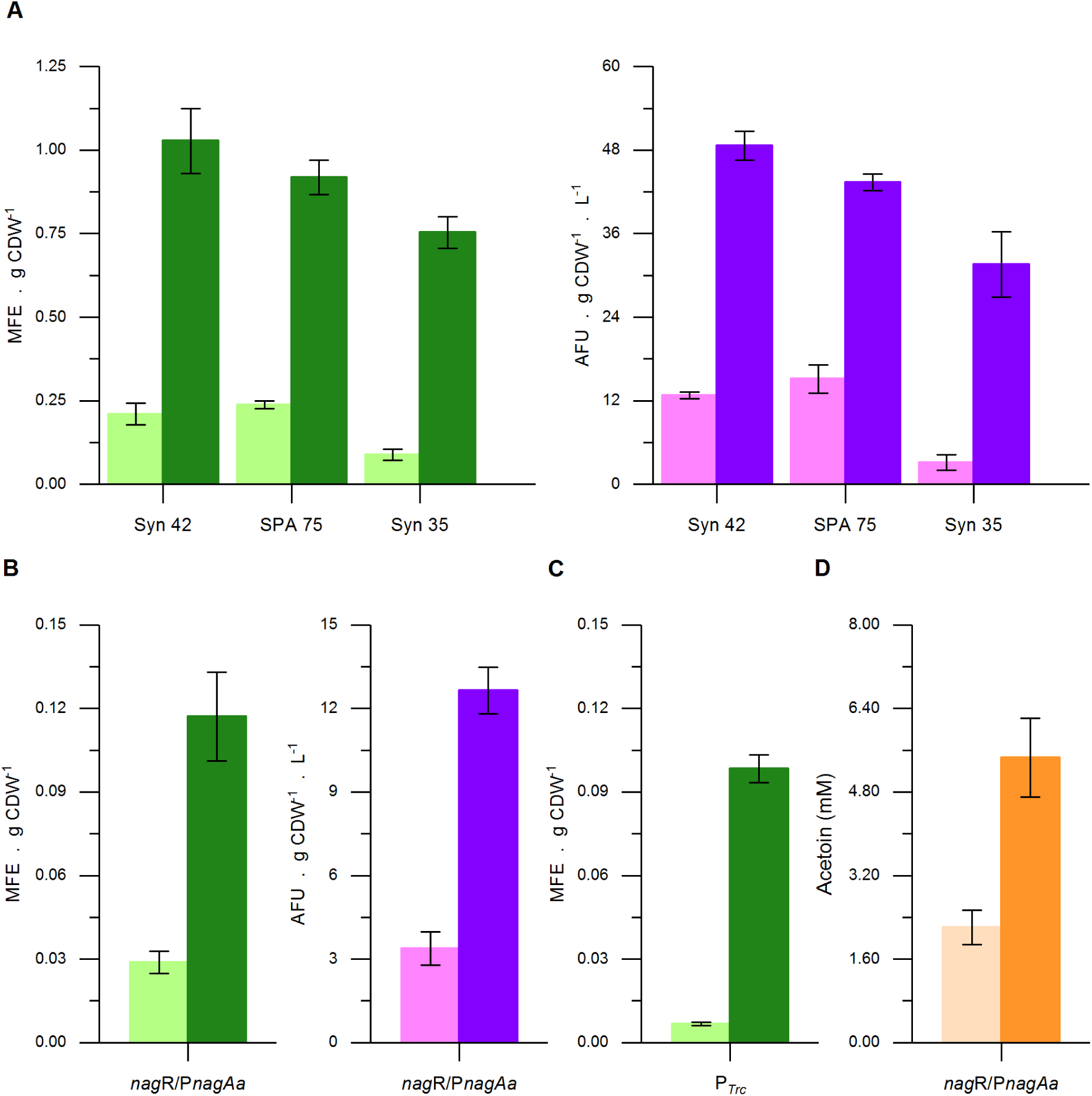
Evaluation of the developed gene expression constructs: plasmid-based expression of msfGFP (green) and mCherry (pink) with the traditional (light-colored) or optimized (dark-colored) expression cassette employing (A) constitutive and (B) the inducible P*nagAa* promoter, (C) mfsGFP expression under the control of the inducible P_Trc_ promoter from single copies of the traditional and optimized gene expression cassettes integrated into the *att*Tn7 site, (D) plasmid-based expression of an acetoin pathway under the control of the inducible P*nagAa* promoter employing a traditional and an optimized expression cassette Error bars indicate the standard deviation of three biological replicates except for the inducible *nag*R/P*nagAa* construct expressing mCherry in which biological duplicates are represented. CDW, cell dry weight. Abbreviations: MFE, μmoles of fluorescein equivalents; AFU, arbitrary fluorescence units.

In a recent study by Clifton et al.(Clifton et al., 2018) fluorescent protein expression in constructs harboring a RiboJ was evaluated using 24 different, constitutive promoters covering a broad spectrum of expression strength. Specific fluorescence values were not reported prohibiting a direct comparison with our results. Based on the absolute GFP fluorescence values, however, no strong correlation between protein expression and promoter strength was observed.

### 3.2 Characterization of plasmid-based, inducible fluorescent protein expression

Controlled gene expression is required, for instance, when the synthesis of a target product harms cell fitness and needs to be decoupled from growth or in genetic circuits (Chen, 2012; Jusiak et al., 2016; Terpe, 2006; Voigt, 2006). We, therefore, chose to further evaluate inducible gene expression by placing the two fluorescent reporter proteins under control of the *nag*R/P*nagAa* promoter, inducible with low concentrations of the relatively cheap inducer salicylate (Hüsken et al., 2001). In both, the traditional and optimized expression constructs, the bidirectional terminators were left out as the first attempt to reduce the overall size of the cassette. The removal of the bidirectional terminators should bear similar effects in both cassettes as they have the identical genetic context. Aside from the removal of the bidirectional terminators and the use of a different promoter, both expression cassettes contained the equal genetic parts used for the evaluation of the constitutive promoters (Figure 2 B). Under the control of the inducible *nag*R/P*nagAa* promoter, the optimized gene expression cassette showed a similar behavior as with the strong constitutive promoters. The specific fluorescence was increased by 4-fold with the optimized gene expression cassette in comparison to their respective traditional counterparts (Figure 3 B). A similar increase was observed for the msfGFP signal under non-induced conditions whereas for the mCherry constructs only a 2-fold increase was observed. The stable increase of the msfGFP signal under both induced and uninduced conditions supports the concept of a post-transcriptional gene expression enhancement. The lower fold ratio in the non-induced conditions observed with the mCherry constructs could be related to analytical inaccuracies of the weaker mCherry signal. Inducible promoters, such as the *nag*R/P*nagAa* promoter system used in this work, tend to have a basal expression which could become problematic for the expression of toxic genes or their use in systems that need to be tightly regulated, *e. g*., genetic circuits. Attempts to achieve non-leaky inducible expression systems have been made, but their number continues to be limited (Horbal and Luzhetskyy, 2016). We propose the use of the optimized gene expression cassette in known non-leaky inducible promoter setups to increase the available expression range in such systems rather than aiming to engineer a novel non-leaky inducible variant. Since the additional parts of the optimized gene expression cassette tend to act on transcribed mRNA only, an optimized gene expression cassette harboring a tight, inducible promoter system could still exhibit the desired non-basal expression in the absence of the inducer and, once induced, reach higher expression levels than the standard counterpart.

### 3.3 Characterization of inducible fluorescence expression of single, genome-integrated constructs

Genomic integration is generally preferred over plasmid-based expression when it comes to the stable construction of cell factories. Integrating the pathways into the genome grants higher genetic stability since it avoids common plasmid-based expression issues such as plasmid segregation and copy number variability (Jahn et al., 2014; Lindmeyer et al., 2015), plasmid replication-related growth impairment (Mi et al., 2016), and antibiotic dependency. However, genomic integration possesses specific disadvantages like generally significantly lower expression levels and a limited number of characterized integration sites (Otto et al., 2019). A common approach to overcome the low expression levels of single genomic integration is to directly or randomly integrate the desired expression cassette at multiple sites. However, the directed multiple integration procedure is laborious and limited by the number of characterized integration sites, whereas random integration requires high-throughput screening. As we had seen a significant increase in expression strength with the optimized cassettes located on a plasmid, we argued that this device might also be valuable to boost the output of genome-integrated constructs. To evaluate if the optimized gene expression cassette could relief the low expression limitation of genomic integrations, single genomic integration cassettes expressing msfGFP were constructed and integrated at the neutral *att*Tn7 site. Further reduction of the optimized gene expression cassette was attempted by removing the RNAse III site. As the RNAse III site does not exert a gene expression enhancement function but rather contributes to its standardization, negative impact on gene expression should not be anticipated with this omission.

To expand the promoter scope further, the standard LacI-repressed P_Trc_ promoter was chosen to drive the gene expression of the genome integrated cassettes. As in the plasmid-based evaluations, the optimized gene expression cassette harbored the RiboJ and BCD2, whereas the traditional version contained the 2^nd^ RBS of the BCD2 (Figure 2 C). Introducing this optimized gene expression cassette in the single genomic *att*Tn7 locus yielded a ca. 15-fold expression increase in comparison to the traditional analog, the highest fold increase observed in this work (Figure 3 C). Even though RNAse III is not the major endoribonuclease responsible for mRNA turnover in bacteria, the enzyme does contribute to mRNA degradation.(Deutscher, 2006) By removing the RNAse III site from the optimized gene expression cassette, an mRNA degradation target was abolished, which might have resulted in an increased half-life of the transcripts and consequently higher expression levels. The single, genome-integrated optimized expression cassette achieved expression levels of 0.098 ± 0.005 μmol fluorescein g^-1^ cell dry weight (CDW), an expression strength in the range of the plasmid-based optimized cassette under the control of the inducible promoter *nag*R*/*P*nagAa* or the constitutive promoter Syn35 within the traditional cassette. Although the use of different promoters does not allow a fair comparison, it is still noteworthy to state that expression levels were achieved with the single genomic integration cassette, which are usually seen for episomally expressed genes.

### 3.4 Evaluation of a recombinant acetoin production pathway employing the optimized expression cassette

Finally, a heterologous acetoin pathway was framed within the pTN1 plasmid backbone to further evaluate the optimized expression cassette in a production context. The acetoin pathway included a C83S mutant of the *E. coli* K-12 MG1655 acetolactate synthase (*ilvB*) and an acetolactate decarboxylase (*aldB*) from *Brevibacillus brevis* (Figure 2 E). The *E. coli ilv*B C83S mutant was chosen due to its 4-fold lower Km and 126% higher Kcat/Km ratio compared with the wild type, making it one of the most efficient enzymes of this class so far characterized (Belenky et al., 2012). The *B. brevis aldB* was selected based on its low Km value (B. Rostgaard et al., 1987). In the polycistronic design, expression of both genes was driven by the inducible *nag*R/P*nagAa* promoter system while each gene was framed with a ribozyme, a bicistronic design version, and an RNAse III site. The acetolactate synthase was framed within the VtmoJ ribozyme gene, the BCD1 bicistronic design, and the RNAse III R0.5 site whereas the acetolactate decarboxylase was surrounded with the RiboJ ribozyme gene, the BCD2 bicistronic design, and the RNAse III R1.1 site (Figure 2 D). The traditional expression cassette was obtained by removing the expression enhancing genetic parts while maintaining the 2^nd^ RBS of BCD1 upstream of the *ilvB* and the 2^nd^ RBS of BCD2 upstream of the *aldB* (Figure 2 D).

Employing the optimized gene expression architecture to the acetoin pathway led to an acetoin accumulation of 5.5 ± 0.76 mM, representing a 2.5-fold production increase when compared to the traditional counterpart (Figure 3 C). The increase of acetoin production solely using the optimized gene expression cassette shows that its use can be extended to production contexts. The optimized gene expression cassette could be advantageous for metabolic engineering approaches where high gene expression is required, such as redirecting native metabolites to the desired production pathway or for enzyme production for *in vitro* applications.

### 3.5 qPCR based elucidation of mRNA stability

The high fluorescence expression levels and increased acetoin production achieved by the optimized gene expression in this work arose from the combination of mRNA stabilizing and translation boosting genetic parts. Lou et al. integrated hairpins in both ribozymes used in this work, RiboJ and VtmoJ, to expose the RBS and confirmed their catalytic functionality through rapid amplification of cDNA 5’-ends.(Lou et al., 2012) The presence of such hairpins in the 5’UTR region was reported to increase the respective mRNA half-life, which leads to higher protein production.(Viegas et al., 2018) To evaluate if such a phenomenon occurred in the optimized gene expression cassette, qPCR assays were performed to compare the decay rates of the transcripts. For this purpose, cultivations of the strains expressing *msfGFP* under the control of the SPA75 promoter were treated with rifampicin and nalidixic acid in the early exponential growth phase, and mRNA samples were retrieved over time. The addition of rifampicin inhibits bacterial RNA polymerase, whereas nalidixic acid inhibits a subunit of the DNA gyrase and topoisomerase. As the mRNA abundance data of the *mfsGFP* were normalized with the data of the housekeeping gene *rpoB* to cancel out small differences in the template concentration, the determination of the absolute decay rate of the *mfsGFP* transcript was not possible. Instead, we calculated the difference between the decay rates of the *rpoB* and the *msfGFP* mRNA (i.e., *k*_*rpoB*_ ™ *k*_*msfGFP*_, with *k* being the decay rate). Given that the *rpoB* mRNA decay should be equal in both strains the delta can be used to reveal a possible difference in the *msfGFP* decay rates in the optimized and traditional expression architecture. As hypothesized, the data was significantly higher for the optimized construct (1.6-fold), which translates to a reduced decay rate of the *msfGFP* transcript (Figure 4).

**Figure 4.**
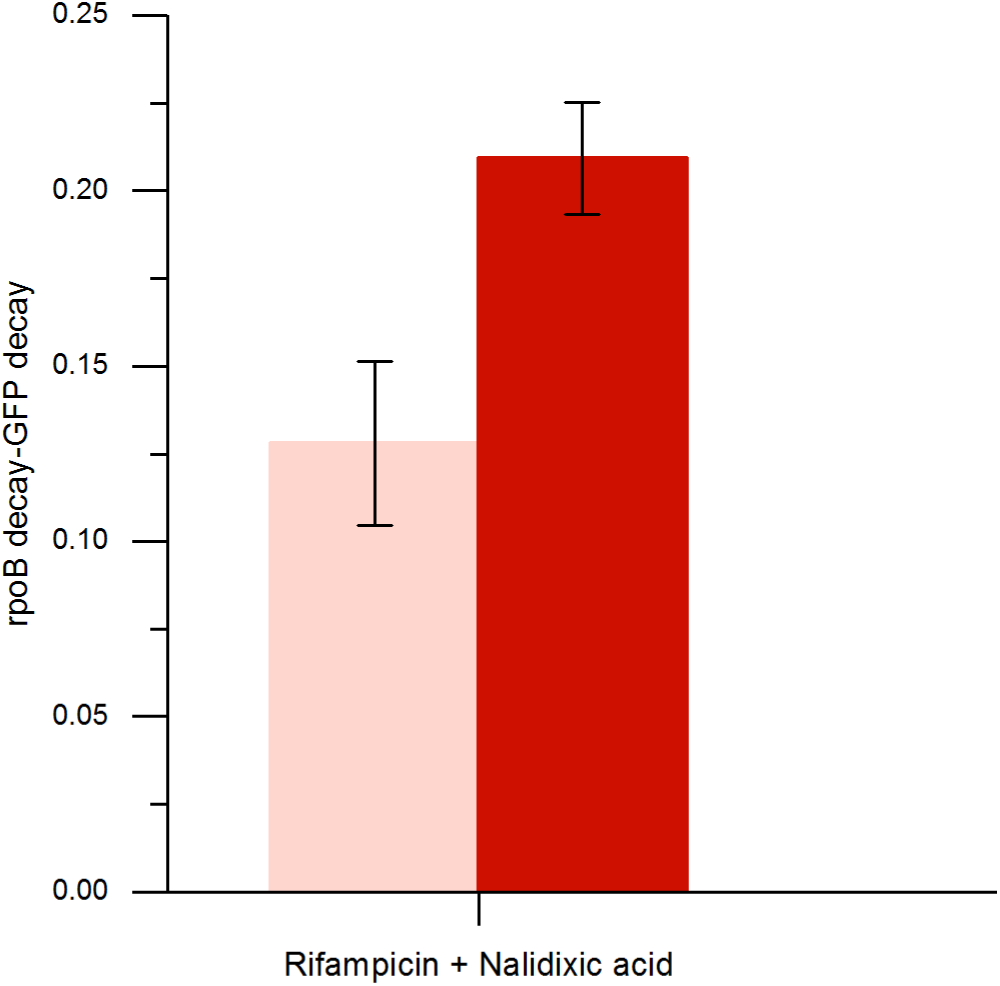
qPCR based elucidation of mRNA stability of *msfGFP* transcripts from the plasmid-based traditional (light-colored) and optimized (dark-colored) expression cassettes harboring the constitutive SPA75 promoter. Early exponential cultivations were treated with the antibiotics, nalidixic acid and rifampicin, to halt DNA replication and transcription, respectively. Samples were taken over time, and mRNA levels were assessed through qPCR. mRNA decay rates of transcripts from each expression cassette were retrieved through the difference between the decay rate of the housekeeping gene *rpoB* and the target *msfGFP*. Error bars indicate the standard deviation of two biological replicates.

Hence, higher mRNA stability is indeed one factor contributing to the observed increase in fluorescence output and acetoin production.

## 4 Conclusion

In this study, the optimized gene expression cassette architecture proposed by Nielsen et al. was evaluated in *P. taiwanensis* VLB120 for the expression of fluorescent reporter genes and a 2-step acetoin biosynthesis operon. The optimized gene expression cassette was characterized on a plasmid or single genomic integration basis with either constitutive or inducible promoters to cover all commonly used expression approaches in metabolic engineering. In all evaluations, the optimized gene expression cassette outperformed its traditional counterpart. The highest improvement fold was observed once the RNAse III site was removed and evaluated on a single genomic integration basis under the control of the IPTG inducible P_*Trc*_ promoter. Such a boost allowed a single genomic integration-based expression to achieve expression levels commonly reached with plasmids. Within the constitutive promoter paradigm, the optimized gene expression cassette increased expression levels of the strongest promoter of a promoter library, showing that this tool could be used to extend expression ranges further. The mRNA transcripts retrieved by the optimized gene expression cassette harnessed higher stability than the transcripts from the traditional counterpart, validating that mRNA stability contributed to the observed results. This work demonstrates the applicability of the optimized gene expression cassette as a tool to achieve high gene expression levels through transcription-independent approaches that rely on mRNA stability and translation efficiency.

## 5 Acknowledgments

We thank Maike Otto and Nick Wierckx for valuable discussions.

## 6 Supplementary material

Primers, annotated sequences of ordered DNA fragments, conversion of Biolector fluorescence a.u. into μM fluorescein and qPCR primer pair efficiencies can be found in the supplementary material.

## 7 Author Contributions

DN and BEE conceived the study with the help of LMB. BEE and LMB supervised the study. DN performed all experiments with the support of SV. All authors analyzed the data. All authors have approved the final version of the manuscript.

