## Supplementary Information for "*Pseudomonas* mRNA 2.0: Boosting Gene Expression Through Enhanced mRNA Stability and Translational Efficiency"

Table S1. Primers used for cloning and qPCR. Lower-case letters represent nucleotide overhangs used for Gibson cloning. Upper-case letters represent nucleotides binding to the amplicon.

| Primer | Sequence | Description |
| --- | --- | --- |
| DN_39 | gcatggatgaactctacaaataATAGAGGGACAAACTCAAGGTC | Opt_Syn42_GFP backbone (fwd) |
| DN_40 | cctttatgaattcccatgaTCATTAGAAAACCTCCTTAGCATG | Opt_Syn42_GFP backbone (rev) |
| DN_41 | CATGCTAAGGAGGTTTTCTAATGATCATGGGAATTCATAAAGG | Opt_Syn42/Syn35_GFP insert (fwd) |
| DN_42 | GACCTTGAGTTTGTCCCTCTATTATTTGTAGAGTTCATCCATGC | Opt_Syn42/Syn35/SPA75_GFP insert (rev) |
| DN_55 | gaactctacaaataaAACGAGAAAAGCCAACCTGCGGGTTGG | Tra_Syn42 backbone (fwd) |
| DN_56 | cattagaaaacctcCTCCTAGGCGTGCAATTATACCTGGCCGC | Tra_Syn42 backbone (rev) |
| DN_57 | taattgcacgcctaggAGGAGGTTTTCTAATGATCATGGG | Tra_Syn42_GFP insert (fwd) |
| DN_58 | gttggcttttctcgttTTATTTGTAGAGTTCATCCATGCCG | Tra_Syn42_GFP insert (rev) |
| DN_247 | AACGAGAAAAGCCAACCTG | Tra_Syn35/SPA75_GFP; Tra_Syn42/Syn35/SPA75_mCherry backbone (fwd) |
| DN_248 | TAGAAAACCTCCTCCTAGGCG | Tra_Syn42_mCherry backbone (rev) |
| DN_270 | CCTAGGCCCCAAATTATAATTCTAAAC | Tra_Syn35_GFP; Tra_Syn35_mCherry backbone (rev) |
| DN_274 | CCTAGGCGTGCAATTATAGTATC | Tra_SPA75_GFP; Tra_SPA75_mCherry backbone (rev) |
| DN_249 | gcctaggaggaggttttctaATGGTGAGCAAGGGCGAG | Tra_Syn42_mCherry insert (fwd) |
| DN_250 | gcaggttggcttttctcgttTTACTTGTACAGCTCGTCCATG | Tra_Syn42/SPA75_mCherry insert (rev) |
| DN_271 | attataatttggggcctaggAGGAGGTTTTCTAATGATC | Tra_Syn35_GFP insert (fwd) |
| DN_272 | gcaggttggcttttctcgttTTATTTGTAGAGTTCATCCATG | Tra_Syn35/SPA75_GFP insert (rev) |
| DN_281 | attataatttggggcctaggAGGAGGTTTTCTAATGGTG | Tra_Syn35_mCherry insert (fwd) |
| DN_250 | gcaggttggcttttctcgttTTACTTGTACAGCTCGTCCATG | Tra_Syn35_mCherry insert (rev) |
| DN_275 | actataattgcacgcctaggAGGAGGTTTTCTAATGATC | Tra_SPA75_GFP insert (fwd) |
| DN_276 | actataattgcacgcctaggAGGAGGTTTTCTAATGGTG | Tra_SPA75_mCherry (fwd) |
| DN_229 | TAGAGGGACAAACTCAAG | Opt_Syn35/SPA75_GFP; Opt_Syn42/Syn35/SPA75_mCherry backbone (fwd) |
| DN_230 | TAGAAAACCTCCTTAGCATG | Opt_Syn35_GFP; Opt_Syn42/Syn35/SPA75_mCherry backbone (rev) |
| DN_267 | TTAATTAAGACGTCTTGACATAAGC | Opt_SPA75_GFP backbone (rev) |
| DN_245 | catgctaaggaggttttctaATGGTGAGCAAGGGCGAG | Opt_Syn42/Syn35/SPA75_mCherry insert (fwd) |
| DN_236 | accttgagtttgtccctctaTTACTTGTACAGCTCGTCCATG | Opt_Syn42/Syn35/SPA75_mCherry insert (rev) |
| DN_269 | tgtcaagacgtcttaattaaGCCCATTGACAACACTATTTTTTGATACTATAATTGCACGCCTAGGAGCTGTCACCGG | Opt_SPA75_GFP insert (fwd) |
| DN_127 | CGGCCGCGCTAGCACTGA | *nag*R_Tra/Opt_GFP/mCherry/ilvB_aldB backbone (fwd) |
| DN_288 | CCGACGTCGCATGCTCCT | *nag*R_Tra/Opt_GFP/mCherry/ilvB_aldB backbone (rev) |
| DN_289 | agaggagcatgcgacgtcggAGGAGGTTTTCTAATGATC | *nag*R_Tra_GFP insert (fwd) |
| DN_142 | ggtcagtgctagcgcggccgTTATTTGTAGAGTTCATCCATGC | *nag*R_Tra_GFP insert (rev) |
| DN_290 | agaggagcatgcgacgtcggAGCTGTCACCGGATGTGC | *nag*R_Opt_GFP/mCherry insert (fwd) |
| DN_291 | ggtcagtgctagcgcggccgAAGAAGGTCAATCATAAAGGCCAC | *nag*R_Opt_GFP/mCherry/*aldB* insert (rev) |
| DN_292 | agaggagcatgcgacgtcggAGGAGGTTTTCTAATGGTG | *nag*R_Tra_mCherry insert (fwd) |
| DN_293 | ggtcagtgctagcgcggccgTTACTTGTACAGCTCGTC | *nag*R_Tra_mCherry insert (rev) |
| DN_298 | agaggagcatgcgacgtcggACAGGAGACTTTCTAATGGCC | *nag*R_Tra_*ilvB* insert (fwd) |
| DN_299 | ctaggcgtgcTCACTCGCCGACCATCTC | *nag*R_Tra_*ilvB* insert (rev) |
| DN_300 | cggcgagtgaGCACGCCTAGGAGGAGGTTTTC | *nag*R_Tra_*aldB* insert (fwd) |
| DN_301 | ggtcagtgctagcgcggccgTCACTTGCGCTCGCTTTC | *nag*R_Tra_*aldB* insert (rev) |
| DN_302 | agaggagcatgcgacgtcggAGTCCGTAGTGGATGTGTATC | *nag*R_Opt_*ilvB* insert (fwd) |
| DN_303 | ctaggcgtgcAAAGTGATAATCATAAAGGCCAC | *nag*R_Opt_*ilvB* insert (rev) |
| DN_294 | ttatcactttGCACGCCTAGGAGCTGTC | *nag*R_Opt_*aldB* insert (fwd) |
| DN_537 | AGGAGGTTTGTATCTCTAATG | *att*Tn7_Tra_GFP backbone (fwd) |
| DN_534 | AGCTGTCACCGGATGTGC | *att*Tn7_Opt_GFP backbone (fwd) |
| DN_415 | TTGACAGCTTATCATCGATAAAC | *att*Tn7_Tra/Opt_GFP backbone (rev) |
| DN_532 | tatcgatgataagctgtcaaTTGACACCATCGAATGGTGC | *att*Tn7_Tra/Opt_GFP insert (fwd) |
| DN_538 | attagagatacaaacctcctGCGGCCTAGGGTGTGAAATTG | *att*Tn7_Tra_GFP insert (rev) |
| DN_539 | aagcacatccggtgacagctGCGGCCTAGGGTGTGAAATTG | *att*Tn7_Opt_GFP insert (rev) |
| DN_392 | CACCGCAGACAAACAGAAGA | qPCR primer *msf*GFP (fwd)(Otto et al., 2019) |
| DN_393 | ACTGGGTGGACAGGTAGTGG | qPCR primer *msf*GFP (rev)(Otto et al., 2019) |
| DN_549 | TTGGCCCAGAGGAAATCAC | qPCR primer rpoB (fwd) |
| DN_550 | GGCACCGACGTAGACAATAC | qPCR primer rpoB (fwd) |

Otto, M., Wynands, B., Drepper, T., Jaeger, K.-E., Thies, S., Loeschcke, A., et al. (2019). Targeting 16S ribosomal DNA for stable recombinant gene expression in Pseudomonas. *ACS Synth. Biol.*, acssynbio.9b00195. doi:10.1021/acssynbio.9b00195.

**Supplementary Information A**

*aldB, B. brevis*

CCATGGTACCACCGTCAAAAAAAACGGCGCTTTTTAGCGCCGTTTTTATTTTTCAACCTTCGCATACGCTACTTGCATTACAGTTTACGAACCGAACAGGCTTATGTCAAGACGTCTTAATTAAGCCCATTGACAAGGCTCTCGCGGCCAGGTATAATTGCACGCCTAGGAGCTGTCACCGGATGTGCTTTCCGGTCTGATGAGTCCGTGAGGACGAAACAGCCTCTACAAATTTTGTTTAAGCCCAAGTTCACTTAAAAAGGAGATCAACAATGAAAGCAATTTTCGTACTGAAACATCTTAATCATGCTAAGGAGGTTTTCTAATGAAGAAGAACATTATCACGTCGATTACCAGCTTGGCGTTGGTCGCGGGCCTCAGCTTGACCGCGTTCGCCGCAACGACCGCCACGGTGCCCGCCCCCCCGGCCAAGCAGGAAAGCAAGCCCGCCGTCGCCGCCAACCCGGCTCCTAAGAATGTGCTGTTCCAGTACAGCACCATCAACGCCCTTATGCTGGGCCAGTTCGAAGGCGACCTGACGTTGAAGGATCTGAAGTTGCGCGGCGATATGGGCCTGGGCACGATCAACGATCTTGACGGCGAAATGATCCAAATGGGCACCAAATTCTACCAAATCGACTCCACGGGCAAACTGAGCGAACTCCCAGAATCCGTAAAGACCCCATTCGCCGTCACGACCCACTTCGAGCCAAAGGAGAAGACGACCCTGACCAACGTGCAGGATTACAACCAGCTGACCAAGATGCTGGAGGAAAAATTCGAGAACAAAAACGTCTTCTACGCCGTAAAACTGACCGGGACCTTCAAGATGGTGAAGGCCCGCACCGTGCCGAAGCAAACCCGTCCATACCCACAACTGACCGAGGTGACGAAGAAGCAGAGCGAGTTCGAGTTCAAGAACGTGAAGGGTACGCTGATCGGCTTCTACACGCCGAACTACGCCGCCGCCCTGAATGTCCCCGGTTTCCATTTGCATTTCATCACGGAGGACAAAACGTCCGGCGGTCATGTACTGAACCTTCAGTTCGATAATGCGAACCTGGAGATCAGCCCCATCCACGAGTTCGACGTCCAGCTGCCCCATACCGATGATTTCGCCCATTCGGATTTGACCCAGGTGACCACGTCGCAAGTACATCAGGCCGAAAGCGAGCGCAAGTGATAGAGGGACAAACTCAAGGTCATTCGCAAGAGTGGCCTTTATGATTGACCTTCTTAACGAGAAAAGCCAACCTGCGGGTTGGCTTTTTTATGCAGCGGCCGC

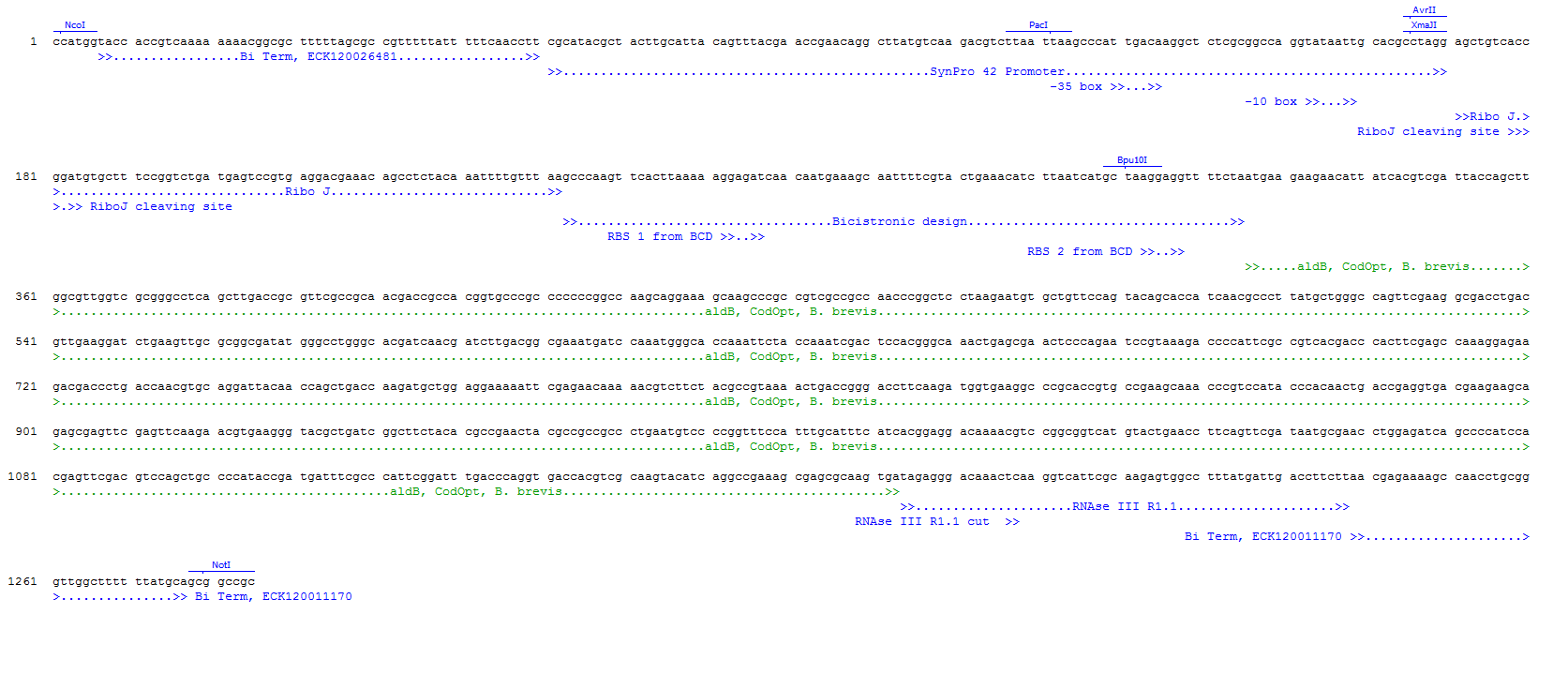

**Figure 1.** Annotated sequence of the synthetic DNA fragment ordered, containing the bidirectional terminators, Syn42 promoter, RiboJ, BCD2, codon optimized *aldB* gene from *B. brevis* and RNAse III site R1.1.

*ilvB* C83S*, E. coli* K12

CCATGGAACGAGAAAAGCCAACCTGCGGGTTGGCTTTTTTATGCACGCATACGCTACTTGCATTACAGTTTACGAACCGAACAGGCTTATGTCAAGACGTCTTAATTAATCTACTTGACATCCGACATTCGCGACTGTATAATAAGTTGACCTAGGGAGTCCGTAGTGGATGTGTATCCACTCTGATGAGTCCGAAAGGACGAAACGGACCTCTACAAATAATTTTGTTTAAGGGCCCAAGTTCACTTAAAAAGGAGATCAACAATGAAAGCAATTTTCGTACTGAAACATCTTAATCATGCACAGGAGACTTTCTAATGGCCAGCAGCGGCACCACCAGCACCCGCAAACGCTTCACGGGCGCCGAGTTCATCGTCCACTTCTTGGAGCAGCAGGGCATCAAGATCGTCACCGGCATCCCTGGCGGCAGCATCCTGCCGGTGTACGATGCCCTCAGCCAGAGCACCCAGATCCGCCACATCCTGGCTCGCCATGAACAAGGCGCGGGCTTCATCGCCCAGGGCATGGCCCGCACCGACGGCAAGCCCGCCGTCTGCATGGCGTCGAGCGGTCCGGGCGCCACCAATCTGGTCACCGCAATCGCCGACGCCCGTTTGGATAGCATCCCGCTGATCTGCATCACGGGCCAGGTGCCAGCCAGCATGATAGGCACCGATGCCTTCCAGGAGGTGGACACCTACGGCATCAGCATCCCCATCACCAAGCATAACTACTTGGTGCGCCACATCGAGGAACTCCCGCAGGTGATGTCCGATGCCTTCCGCATCGCCCAGTCGGGTCGGCCAGGCCCAGTTTGGATCGATATCCCGAAAGACGTCCAGACCGCCGTGTTCGAAATCGAAACCCAGCCCGCGATGGCTGAGAAAGCCGCGGCTCCGGCCTTCAGCGAAGAAAGCATCCGCGACGCCGCGGCTATGATCAACGCCGCAAAGCGCCCCGTGCTGTACCTGGGCGGCGGTGTCATCAATGCCCCAGCACGCGTGCGCGAACTGGCCGAGAAGGCCCAGCTTCCGACCACCATGACCCTTATGGCTCTGGGTATGCTGCCGAAGGCTCACCCGCTCTCGCTGGGTATGCTCGGGATGCACGGCGTCCGGAGCACCAACTACATCCTCCAGGAGGCCGACCTGCTGATCGTCCTGGGCGCCCGCTTCGACGACCGTGCCATCGGCAAAACCGAGCAGTTCTGCCCGAACGCCAAAATCATCCATGTTGACATTGACCGCGCGGAGTTGGGCAAGATCAAGCAGCCGCACGTGGCCATCCAGGCGGATGTGGACGACGTGCTGGCCCAGCTCATCCCGCTCGTGGAGGCACAGCCGCGCGCCGAATGGCACCAGCTGGTGGCGGACCTTCAACGCGAGTTCCCTTGCCCCATCCCCAAGGCCTGCGATCCCCTGAGCCATTACGGTCTGATCAACGCTGTGGCCGCGTGCGTCGATGACAACGCGATCATCACCACCGATGTGGGTCAACACCAGATGTGGACCGCTCAGGCGTACCCGCTGAACCGCCCGCGCCAGTGGCTCACCAGCGGCGGCCTGGGCACGATGGGGTTCGGTCTGCCCGCGGCCATCGGGGCTGCCCTGGCTAACCCAGACCGCAAGGTGCTGTGCTTCAGCGGTGACGGGAGCCTGATGATGAACATCCAGGAGATGGCCACCGCCAGCGAGAACCAGCTCGACGTCAAGATCATTCTGATGAACAACGAAGCCCTGGGCTTGGTACACCAGCAGCAGAGCCTGTTCTATGAACAGGGCGTCTTCGCCGCAACCTACCCCGGCAAGATTAACTTCATGCAGATCGCAGCCGGGTTCGGGCTGGAAACCTGCGATCTCAATAATGAGGCTGACCCGCAGGCGTCGCTCCAGGAAATCATCAACCGGCCCGGCCCGGCCCTGATCCATGTCCGTATCGACGCCGAGGAGAAGGTGTATCCAATGGTGCCCCCCGGCGCCGCCAACACGGAGATGGTCGGCGAGTGAAGTGATAGACTCAAGGTCGCTCCTAGCGAGTGGCCTTTATGATTATCACTTTAAATAAAAAAGGCACGTCAGATGACGTGCCTTTTTTCTTGTGCGGCCGC

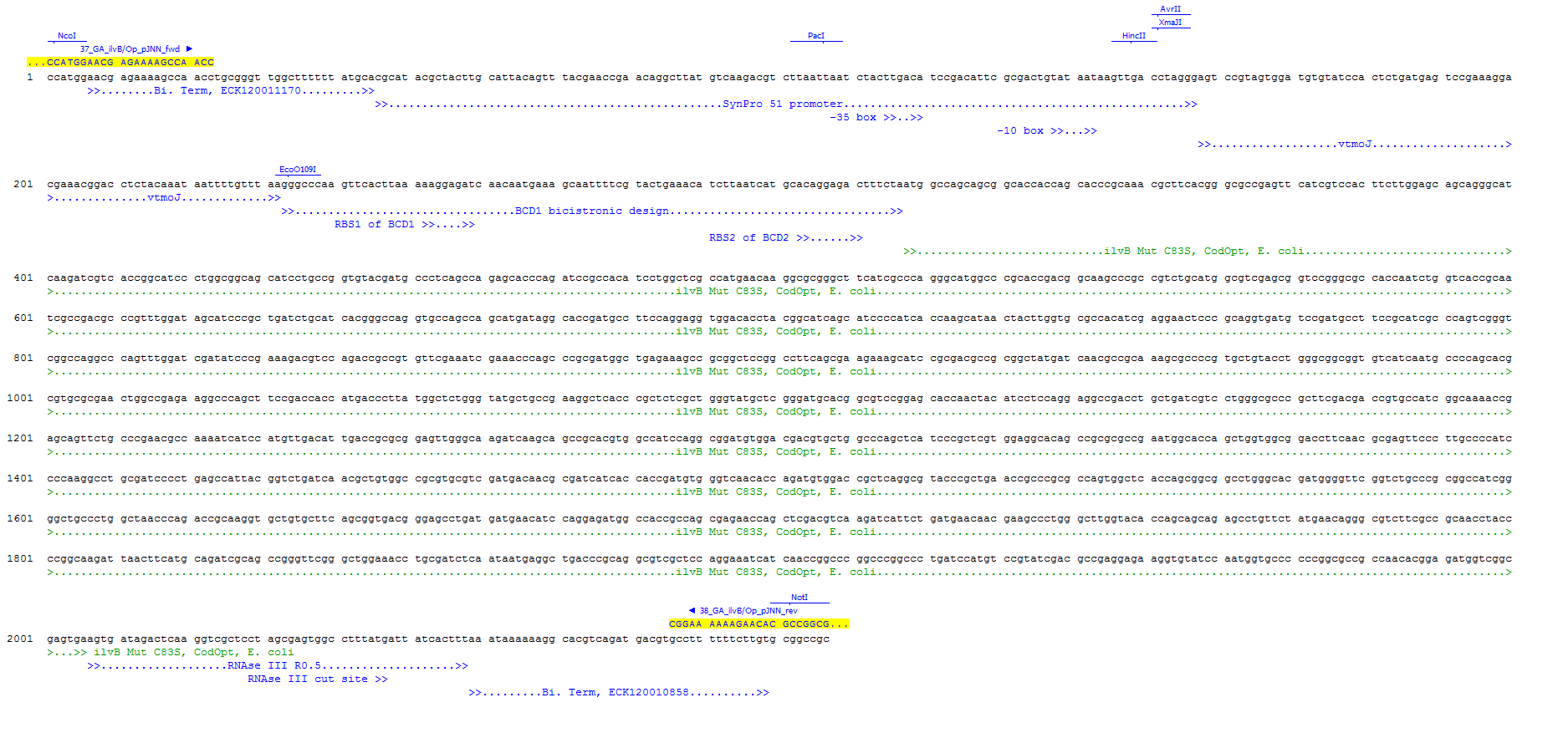

**Figure 2.** Annotated sequence of the synthetic DNA fragment ordered, containing the bidirectional terminators, Syn51 promoter, VtmoJ, BCD1, codon optimized *ilvB* C83S gene from *E. coli* and RNAse III site R0.5.

**Supplementary Information B**

Fluorescein in 0.1mM Borate buffer pH 9.4 (uM)

| a.U. Fluorescence Biolector (488nm/520nm) | Gain 50 |
| --- | --- |
|  | 50 |
|  | 25 |
|  | 12.5 |
|  | 6.25 |
| 1063.72 | 3.125 |
| 590.251 | 1.56 |
| 311.706 | 0.781 |
| 159.619 | 0.39 |
| 80.8723 | 0.195 |
| 41.0483 | 0.0976 |
| 20.553 | 0.0488 |
| 0.063 | 0 |

| a.U. Fluorescence Biolector (488nm/520nm) | Gain 60 |
| --- | --- |
|  | 50 |
|  | 25 |
|  | 12.5 |
|  | 6.25 |
|  | 3.125 |
| 1308.69 | 1.56 |
| 690.175 | 0.781 |
| 354.036 | 0.39 |
| 179.283 | 0.195 |
| 91.023 | 0.0976 |
| 45.6577 | 0.0488 |
| 0.130333 | 0 |

| a.U. Fluorescence Biolector (488nm/520nm) | Gain 70 |
| --- | --- |
|  | 50 |
|  | 25 |
|  | 12.5 |
|  | 6.25 |
|  | 3.125 |
|  | 1.56 |
| 1425.99 | 0.781 |
| 731.644 | 0.39 |
| 370.545 | 0.195 |
| 187.91 | 0.0976 |
| 94.6513 | 0.0488 |
| 0.309333 | 0 |

**Figure 3.** Linear regressions of standards of fluorescein in 0.1 mM borate buffer at pH 9.4 and respective fluorescence a.u. measured with the Biolector set at different gains.

**Supplementary Information C**

CT (Threshold cycle)

| Log [plasmid DNA] | GFP |
| --- | --- |
|  | 11.1233 |
|  | 14.2867 |
| 0 | 17.9533 |
| -1 | 21.89 |
| -2 | 25.12 |
| -3 | 28.6267 |

| Slope | -3.525 |
| --- | --- |
| Efficiency | 0.92171 |

**Figure 4.** qPCR primer pair efficiency for the target gene *msfGFP*.

rpoB

CT (Threshold cycle)

| Log [genomic DNA] | ilvB |
| --- | --- |
| 1 | 15.37 |
| 0 | 18.72 |
| -1 | 21.98 |
| -2 | 25.25 |
| -3 | 28.54 |

| Slope | -3.287 |
| --- | --- |
| Efficiency | 1.014785 |

**Figure 5.** qPCR primer pair efficiency for the housekeeping gene *rpoB*.
